# Identification of genes with oscillatory expression in glioblastoma – The paradigm of *SOX2*

**DOI:** 10.1101/2023.11.12.566579

**Authors:** R.Z Fu, O Cottrell, L Cutillo, A Rowntree, Z Zador, H Wurdak, N Papalopulu, E. Marinopoulou

## Abstract

Quiescence, a reversible state of cell-cycle arrest, is an important state during both normal development and cancer progression. For example, in glioblastoma (GBM) quiescent glioblastoma stem cells (GSCs) play an important role in re-establishing the tumour, leading to relapse. While most studies have focused on identifying differentially expressed genes between proliferative and quiescent cells as potential drivers of this transition, recent studies have shown the importance of protein oscillations in controlling the exit from quiescence of neural stem cells. Here, we have undertaken a genome-wide bioinformatic inference approach to identify genes whose expression oscillates and which may be good candidates for controlling the transition to and from the quiescent cell state in GBM. Our analysis identified, among others, a list of important transcription regulators as potential oscillators, including the stemness gene *SOX2*, which we verified to oscillate in quiescent GSCs. These findings expand on the way we think about gene regulation and introduce new candidate genes as key regulators of quiescence.

## Introduction

Quiescence is a state of cell-cycle arrest which, unlike senescence or terminal differentiation, is reversible upon receiving appropriate signals from the environment^1^. Quiescence is thought to be a property commonly found in stem cells and it is important for homeostasis and regeneration by maintaining a pool of cells from which new cells can be produced, upon exit from quiescence, to replenish cells depleted through ageing or injury ^2^. Quiescence is also important in a cancer context; here, it is thought that transition of cancer stem cells to a quiescent state creates a population with the potential to enter the cell cycle at a later time point, thus, contributing to reactivation of a tumour and clinical relapse ^3,4^. The magnitude of this problem is underscored by recent studies suggesting that cancer treatment may push some surviving cancer cells into quiescence from where they can be reactivated ^5,6^.

Reactivation from quiescence may be problematic for any cancer, but it may be particularly an issue for cancers with a high rate of relapse such as glioblastoma. Glioblastoma, also historically referred to as glioblastoma multiforme (GBM), is the most common and most deadly adult primary brain cancer, comprising more than half of all gliomas ^7^. Despite decades of dedicated research and numerous clinical trials collectively covering a range of often innovative treatment modalities, tragically, there remains no cure and relapse is often inevitable ^8^. Thus, there is a great need for innovative scientific approaches to better understand the basic biology of this cancer and in particular, to understand the mechanisms by which cells enter and exit quiescence.

Efforts to identify molecular drivers of quiescence often focus on differential gene expression between normally dividing cells and their dormant counterparts ^1^. However, even with state-of-the-art molecular methods, such differential gene expression, that is causally related to the acquisition of the quiescent state, has proved elusive to define. Thus, some recent studies have focused on epigenetic changes. Indeed, it was shown that in GBMs there is a failure to lock cells into a differentiated state ^9^, creating a state of increased plasticity whereby cells can transition into undesirable states.

In our own recent work, we have provided evidence for a dynamic conceptual framework of controlling the transition to and from quiescence ^10^. Working with normal, i.e. non-cancerous neural stem cells, we have discovered that protein expression oscillations of the mammalian Hairy and Enhancer of Split 1 (HES1) transcription factor (TF), are needed for cells to be able to exit the quiescent state ^10^. Such oscillations have a periodicity, which is typically shorter than the well-known circadian periodicity, and are increasingly recognised as playing an important role in cell-state transitions in normal development, in a number of different models and contexts e.g.^11–13^.

There is emerging evidence that oscillations are also important in a cancer context. Specifically, HES1 has been shown to oscillate in a breast cancer model ^14^, and so have p53 ^15^ and p21 ^16^. In breast cancer cells, HES1 oscillations are important for efficient proliferation ^14^, while p53 oscillations determine the response to radiotherapy ^17,18^. These findings present a new way of understanding cell-state transitions, based on dynamic gene expression that includes entry to and exit from quiescence. But how common are oscillations and which genes are likely to oscillate? Our previous work has shown that the network motif that commonly underlies oscillatory gene expression (autoregulation and microRNA interaction) is very widespread among mammalian TFs^19^. However, because the presence of oscillations is context specific, we argued that a more direct approach was needed for GBM.

Thus, to answer this question, we leveraged two recent developments. First, we took advantage of a recently developed bioinformatic pipeline, which is designed to uncover oscillatory gene expression in single-cell RNA-seq (scRNA-seq) data, irrespective of their periodicity ^20^. This pipeline was developed by Leng et al. ^20^, and optimised by us ^21^ to improve on the statistical robustness and sensitivity of the method. Second, we took advantage of the availability of a plethora of scRNA-seq data from different human GBM tumours ^22^.

By applying this bioinformatic methodology we uncovered up to 3K genes with the potential to oscillate in GBM. We also found that oscillators that are shared across GBM tumours associate with processes that contribute to cancer pathogenesis, highlighting a potential functional role. To validate our findings, we created an endogenous reporter knock-in GBM line for *SOX2* and we optimised a GBM quiescence protocol. We report that *SOX2* oscillates in both proliferative and quiescent stem cells, with a period of 17h.

Our findings provide a global view of oscillatory gene expression in GBM, which shows that it is more common than previously appreciated. They also uncover a previously unknown level of *SOX2* regulation, which may be important for regulating the transition to quiescence.

## Results

### Inference of oscillatory gene expression in GBM tumours

To identify putative oscillators in GBM we interrogated publicly available scRNA-seq data from the Neftel et al. 2019 study, an in-depth transcriptomic study of GBM tumours ^22^. Five tumours (MGH124, MGH125, MGH102, MGH143, MGH115), representing the most common subtypes of GBM tumours in patients, were selected for inference analysis of oscillators. The tumours correspond to neural-progenitor-like (NPC-like) (MGH124, MGH125, MGH102), astrocyte-like (AC-like) (MGH143) and a mixture of astrocyte-like (AC-like) and mesenchymal-like (MES-like) (MGH115) subtypes according to the classification described in Neftel et al. ^22^. To specifically identify important oscillators for tumour formation and progression, we sought first to determine the neoplastic (tumour cell) compartment of each tumour by using the “Label Transfer” method ^23^. (Fig. 1A). “Label Transfer” is a reference-based integrative analysis approach used to transfer previously identified cell identities onto a query dataset ^23^. In our case, we treated the scRNA-seq data from the 5 tumours as the query dataset and used as a reference dataset the cell identities of neoplastic and non-neoplastic (immune, oligodendrocyte precursors cell (OPC), vascular, neuronal, oligodendrocyte and astrocyte) cells in GBM, as previously described in Darmanis et al., 2017 ^24^, to separate the groups of cells with tumour-cell characteristics from their more differentiated or immune infiltrating counterparts (Fig. 1A).

**Figure 1:**
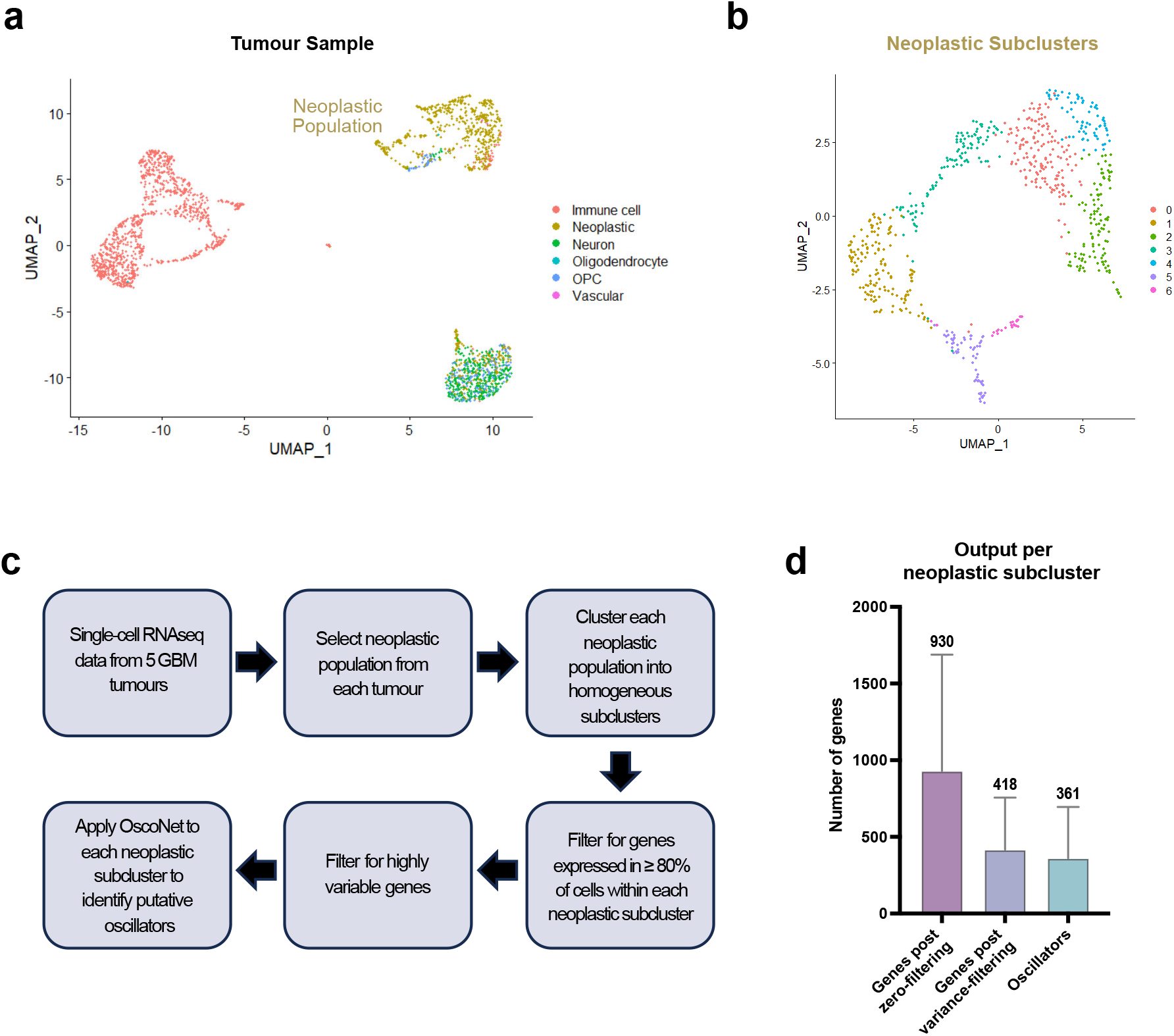
Applying the OscoNet algorithm to identify oscillatory genes in GBM tumours. **(A)** Example UMAP (Uniform Manifold Approximation and Projection) plot representing all single cells in tumour MGH124. Cells belonging to different clusters (neoplastic and non-neoplastic) are identified by different colours **(B)** Example UMAP plot of the neoplastic population of tumour MGH124 showing the clustering of the cells into 7 neoplastic subclusters based on differential gene expression. Different clusters are identified by different colours. **(C)** Diagram showing the sequential steps followed to process the tumour scRNA-seq data prior to subjecting them to inference analysis for oscillatory gene expression using OscoNet. **(D)** Bar graph showing the average number of genes across all neoplastic subclusters across all tumours for each of the pre-processing steps prior to inference analysis for oscillatory gene expression and following inference analysis with OscoNet. Error bars represent standard deviation.

Oscillatory genes are expected to have variable levels between cells thus, we wanted to ensure that we minimise false positive results due to variability in gene expression originating from the mixing of different cell types or states within each neoplastic population. Therefore, additional clustering was performed to identify more homogeneous neoplastic subclusters based on differential gene expression (Fig. 1B). These subclusters revealed appreciable differences in terms of their enrichment for the Human Molecular Signature Database (MSigDB) Hallmark gene sets (Supplementary Table S1).

To infer oscillatory gene expression in the neoplastic populations we used the OscoNet computational algorithm ^21^, an optimised version of a previously described gene oscillatory inference pipeline ^20^. OscoNet capitalises on the fact that within an unsynchronised but homogeneous population, genes which are dynamically expressed are expected to have variable levels that follow a sinusoidal process. To distinguish from noisy expression, and acknowledging that the expression values of genes oscillating with similar frequencies would form an ellipse on a scatter plot (independently of cell order), the algorithm searches for pairs of genes whose expression best fits a two-dimensional sinusoidal function, to uncover genes that oscillate with similar frequencies but possibly different phases ^20^.

OscoNet was applied in each of the neoplastic subclusters separately (Fig. 1B), using the following two filtering criteria. First, for a gene to be considered for OscoNet analysis it had to be expressed in at least 80% of the cells within each subcluster (zero-filtering criterion), as an in-silico exploration determined that the presence of zero expression values affects negatively the performance of the algorithm (Supplementary Methods). Second, genes that had passed the zero-filtering were further filtered based on gene expression variance (variance-filtering criterion), such that only genes whose expression variance was higher than the mean variance of all genes were chosen as an input for the OscoNet pipeline (Fig. 1C). Our analysis revealed that out of all genes that are expressed across each subcluster in ≥ 80% of cells (average of 930 genes), approximately 45% them were found to be highly variable (average of 418 genes) with the majority (87%) of these subsequently inferred to be oscillators (average of 361 genes) (Fig. 1D). Considering that only a handful of genes in the literature have been shown to oscillate to date, this in-silico study suggests that a considerably large number of genes, at least in the setting of GBM, oscillate.

### OscoNet identifies potential oscillators in GBM

The OscoNet inference pipeline identified 361 genes on average to oscillate per neoplastic subcluster per GBM tumour (Fig. 1D), whereas all inferred oscillators across all neoplastic subclusters in all tumours were found to be 3422 (named “All Oscillators”) (Supplementary Table S2). Amongst these, there were known genes that have been previously shown to exhibit oscillatory expression at an ultradian scale (i.e. oscillate with a period less than 24h) and be important for normal neural development. Such genes were HES1, HES4, HES6, Ascl1 and Olig2 (Supplementary Table S2) ^11,25,26^. In addition, we found that cell cycle regulators (GO_0007049, selected those expressed in ≥ 80% of cells in at least one neoplastic subcluster) (Supplementary Table S3), whose expression may oscillate during the cell cycle ^27^, were significantly enriched in our predicted list of oscillators across all tumours (Fig. 2A). These findings suggest that the OscoNet pipeline can successfully identify known oscillators.

**Figure 2:**
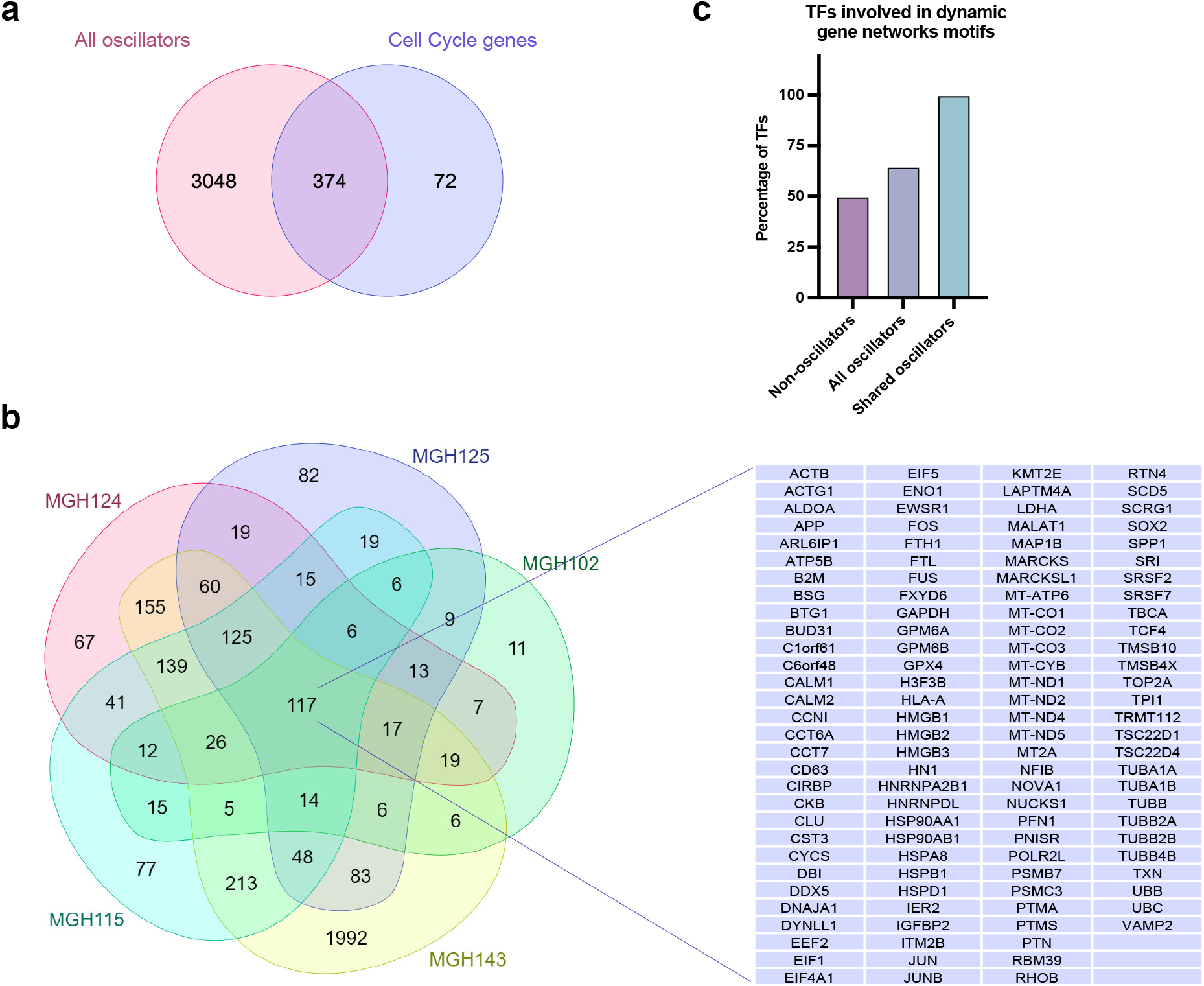
Inferred oscillators enrich for cell cycle genes and are involved in dynamic gene network motifs. **(A)** Venn diagram showing the overlap of “All oscillators” from all 5 tumours (3422 genes) with cell cycle regulators (expressed in at least one tumour) (446 genes). The hypergeometric probability was estimated to be 0.047. **(B)** Venn diagram showing the overlap of all predicted oscillators in each tumour and across all 5 tumours. Table shows the names of 117 oscillators identified to be shared across all tumours. **(C)** Bar graph showing the percentage of TFs in the “Non-oscillators”, “All oscillators” and “Shared oscillators” lists that are involved in a dynamic gene network motif.

To independently verify our predictions, we cross-referenced our findings with that of an alternative bioinformatic screen for genes capable of dynamic gene expression, including oscillations. We have previously found that transcription regulators with the potential to autoregulate (i.e. bind to their own promoter) and which are involved in dual feedback interactions with a miRNA are very common in mammalian TFs ^19^. In particular, in humans, we identified 582 TFs to potentially autoregulate and be in a feedback loop with a miRNA (we call this a “dynamic gene network motif”) ^19^ (Supplementary Table S4). Thus, we sought to identify if any of these genes have been predicted to oscillate by OscoNet in GBM.

We focused on comparing genes with TF activity as the information about genes involved in dynamic gene networks is available only for transcription regulators. We hypothesised that TFs that were inferred to oscillate in GBM were more likely to be involved in such a network as opposed to “Non-oscillators”. In this case, “Non-oscillators” were defined as genes that were expressed in at least one tumour (i.e. passed zero-filtering) but had not been found to oscillate in any of the tumours (643 genes) (named “Non-oscillators”) (Supplementary Table S5). We also hypothesised, that TFs involved in dynamic gene networks would be more prevalent amongst inferred oscillatory TFs that were shared across the 5 tumours, as they were always predicted to oscillate. For the latter, we intersected the oscillators from all neoplastic subclusters per tumour and found a total of 117 oscillatory genes in common (named “Shared oscillators”) (Supplementary Table S6) (Fig. 2B). We then selected only the TFs from each of “All oscillators” (3422), “Shared oscillators” (117) and “Non-oscillators” (643) lists by cross-referencing them against the list of all human TFs as described by Lambert et al., ^28^ (Supplementary Table S7). From these TFs we further selected only those for which it was feasible to assess their involvement in a dynamic gene network motif (i.e. availability of information on autoregulation) and compared them against all TFs that belonged in a dynamic gene network motif (582 genes) (Supplementary Table S4). Importantly, we found that all TFs in the “Shared oscillators” (100%) were involved in network motifs predicted to oscillate as opposed to 64.8% and 50% in the lists of “All oscillators” and “Non-oscillators” respectively (Fig. 2C). These findings lend further support to our method for identifying putative oscillators and suggest that genes predicted to oscillate in every GBM tumour are potentially the highest confidence candidate genes to have dynamic behaviour.

### Oscillatory genes in GBM tumours associate with processes that contribute to cancer pathogenesis

Next, we aimed to gain some insights into the type of genes we have identified that may oscillate in the GBM tumours. As previously mentioned, we considered the “Shared oscillators” to be high confidence oscillators and potentially functionally important. Gene ontology analysis on the “Shared oscillators” revealed that the most enriched molecular and cellular functions were related to cell proliferation and development, cell death and survival, and cell movement, which align with the cancerous profile of the neoplastic population (Fig. 3A). On the contrary, the “Non-oscillators” had lower enrichment scores overall and associated mainly with processes such as protein/molecule trafficking, cell function and maintenance and cellular assembly and organisation (Fig. 3B).

**Figure 3:**
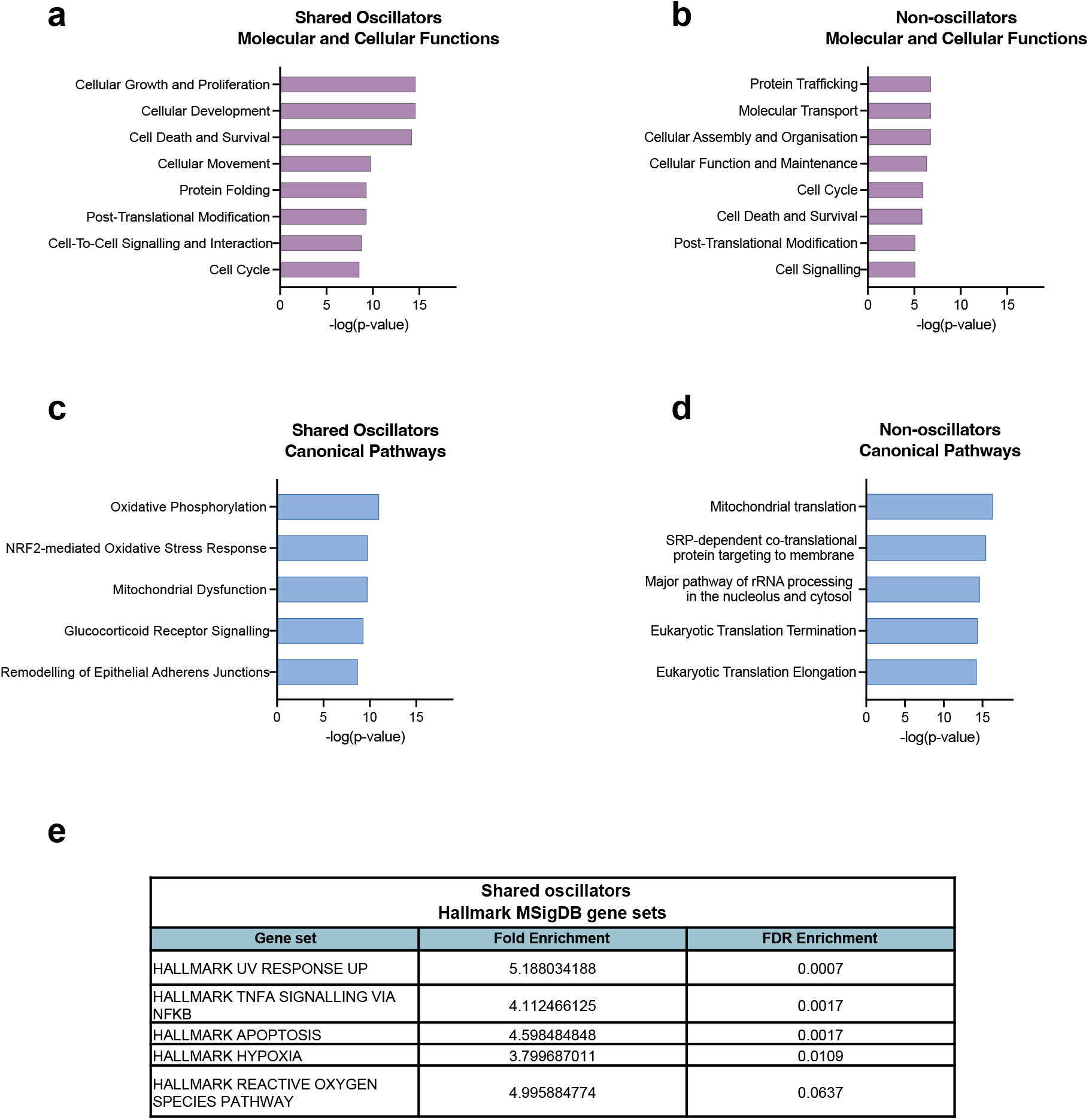
Shared oscillators across the GBM tumours enrich for processes that contribute to cancer pathogenesis. **(A-B)** Bar graphs showing significantly enriched molecular and cellular functions in the lists of “Shared oscillators” (117 genes) **(A)** and “Non-oscillators” (643 genes) **(B)**. **(C-D)** Bar graphs showing significantly enriched canonical pathways in the lists of “Shared oscillators” **(C)** and “Non-oscillators” **(D)**. **(E)** Table showing the fold enrichment and FDR enrichment of “Shared oscillators” for MSigDB Hallmark gene sets. Fold enrichment refers to the percentage of genes in the “Shared oscillators” list that belong in each gene set, divided by the corresponding percentage in the background list. The FDR enrichment reports the hypergeometric test value.

Accordingly, the top enriched canonical pathways in the “Shared oscillators” associated with oxidative stress, mitochondrial dysfunction and remodelling of epithelial adherens junctions, all known to associate with cancer ^29,30^ (Fig. 3C), whereas “Non-oscillators” enriched for core cell processes such as translation and rRNA processing (Fig. 3D).

“Shared oscillators” were also enriched for Hallmark gene sets that are included in MSigDB ^31^. Amongst the most enriched were gene sets that associate with apoptosis, hypoxia and the reactive oxygen species pathway, which all constitute hallmark pathways of cancer ^32^ (Fig. 3E). Conversely, “Non-oscillators” did not enrich for any of the Hallmark gene sets. These findings consistently reveal that oscillators appear enriched in known gene pathways and processes that contribute to cancer pathogenicity whereas “Non-oscillators” tend to associate with more house-keeping processes.

### Oscillatory genes in quiescent/low-cycling GBM tumour cells

Considering that the inevitable tumour relapse is one of the main reasons GBM is practically an incurable disease, we next sought to identify oscillators that are potentially important for driving this process. Recent evidence suggest that the main cause of relapse is the activation of quiescent GBM stem cells (GSCs) that are left behind after surgery, are refractory to chemotherapy and radiotherapy and are able to re-initiate tumour formation ^33^. To this end, we decided to look at oscillators that specifically overlap with a quiescent gene signature or are expressed in the quiescent/low cell-cycling population of the neoplastic cells.

To address this, we interrogated the list of “Shared oscillators” (which we found to be our highest confidence candidates) and particularly the TFs in this list as we argued that an oscillator is more likely to drive cell-state transitions (e.g. exit from quiescence) if they control the expression of downstream genes. We therefore intersected the inferred oscillatory TFs that are shared across the tumours, with a list of GBM neoplastic cell-specific Neural G0 marker genes ^34^ (Supplementary Table S8) (Fig. 4A). The latter had been previously derived by applying a cell-cycle classifier to identify a putative quiescent-like state in gliomas along with the pathways that associate with it ^34^. The overlap of all these lists provided us with a short list of 6 genes (*SOX2, JUNB, FOS, NFIB, TCF4, TSC22D1*) which are predicted to oscillate, regulate transcription and associate with a quiescent phenotype (Fig. 4A).

**Figure 4:**
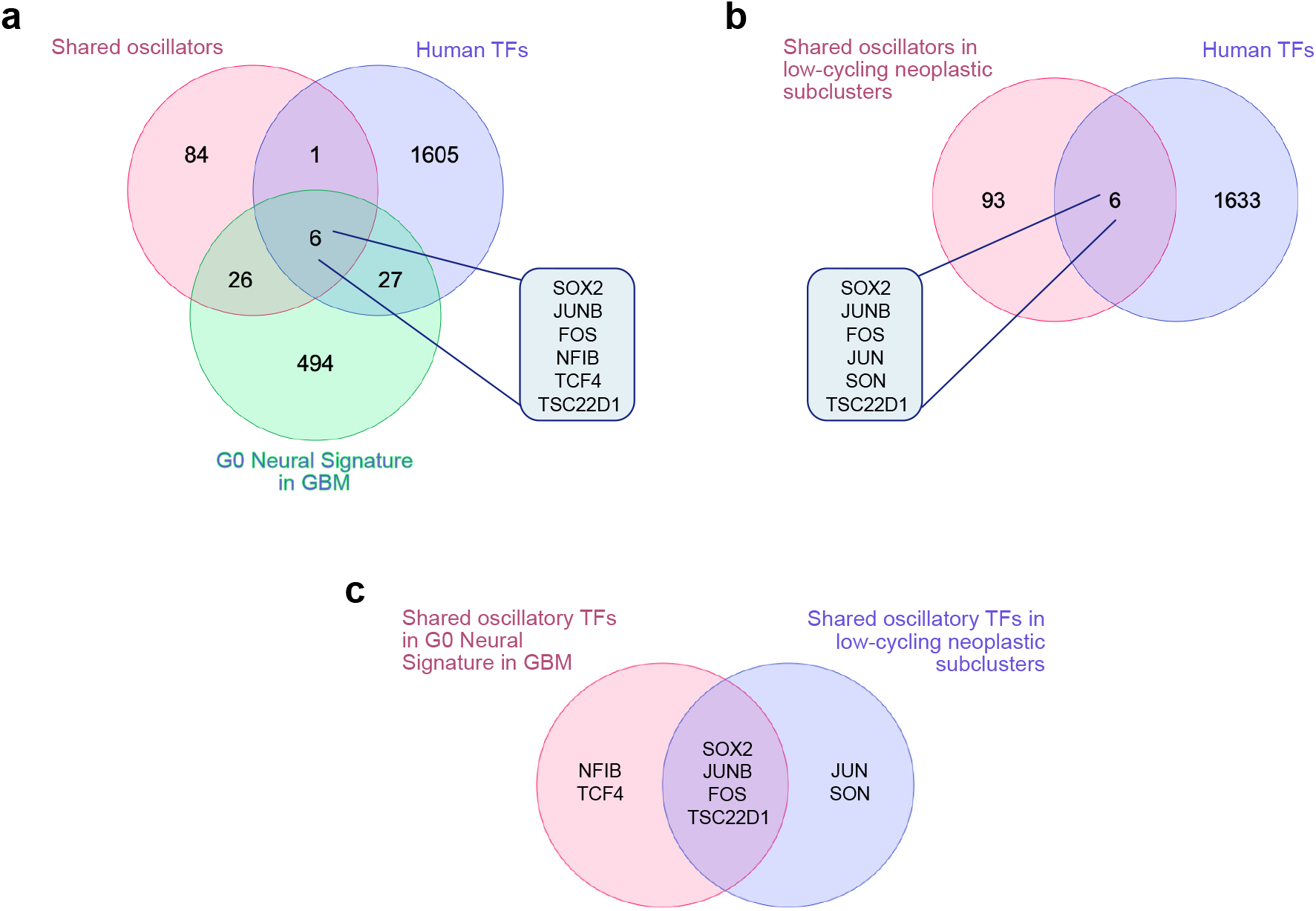
Identification of oscillatory TFs in quiescent/low-cycling GBM tumour cells. **(A)** Venn diagram showing the overlap of “Shared oscillators” (117 genes) with the list of all human TFs (1639 genes) and with the Neural G0 Signature list (553 genes). 6 genes (shown in the table) where identified as common across all three lists. **(B)** Venn diagram showing the overlap of “Shared oscillators” in low-cycling neoplastic subclusters (99 genes) with the list of all human TFs (1639 genes). 6 genes (shown in the table) where identified as common between the two lists. **(C)** Venn diagram showing the overlap of the shared oscillatory TFs in Neural G0 signature with the shared oscillatory TFs in low-cycling neoplastic subclusters.

As an independent way to identify oscillators that potentially control the quiescent state of GBM, we looked at oscillators that were specifically expressed in the low-cell cycling neoplastic subclusters of these tumours, defined as those enriched for Hallmark gene sets associated with the downregulation of cell cycle genes. This analysis identified a list of 99 oscillators shared across the low-cycling neoplastic subclusters in the 5 tumours (Supplementary Table S9) out of which only 6 were found to be putative TFs (Fig. 4B). Interestingly, 4 out those 6 genes (*SOX2, JUNB, FOS, TSC22D1*) were also identified by our previous analysis on the Neural G0 signature-associated oscillators (Fig. 4A) revealing a core of 4 TFs that oscillate and are potentially important for regulating the quiescent state (Fig. 4C).

### Establishing an *in vitro* system to study oscillatory gene expression in quiescence

Having identified a core list of candidate oscillators in low-cycling neoplastic cells, our next objective was to establish a GBM *in vitro* system to verify protein expression oscillations in a quiescent state. Our oscillator inference analysis was based on 5 tumours represented primarily by the NPC-like and MES-like subtypes (proneural and mesenchymal subtype respectively according to Verhaak classification ^35^), with the majority belonging to the NPC-like subtype ^22^. We therefore chose the previously characterised GBM1 patient-derived GSC line as our experimental model, which was found to primarily enrich for a proneural transcriptional signature ^36,37^.

Next, we used the GBM1 cells to induce a quiescent state *in vitro* by replacing the EGF and FGF mitogens in the culture with BMP4. Previous studies have shown that GSCs undergo cell-cycle arrest in response to BMP4, and although they exhibit astrocyte differentiation ^9,37,38^ they subsequently fail to permanently exit the cell cycle and remain susceptible to re-entry ^9^. Accordingly, live image-based confluency analysis of GBM1 cells over a period of 14 days showed that cells amplify over time in the presence of mitogens but they stop within 48h of BMP4 administration and remain cytostatic for as long as they are treated with BMP4 (up to 7 days) (Fig. 5A). Importantly, upon withdrawal of BMP4 and re-introduction of FGF and EGF mitogens in the culture, GBM1 cells were able to re-enter the cell cycle (Fig. 5A). These results suggest that administration and subsequent removal of BMP4 facilitates a reversible cell-cycle arrest suggesting that addition of BMP4 induces a quiescent state. Immunostaining of the cells for the Ki67 proliferation marker in proliferative, quiescent and re-activation conditions confirmed the loss of Ki67 in the presence of BMP4 and re-expression of the marker following re-activation (Fig. 5B-C). Altogether these findings suggest that the GBM1 cell line is a suitable experimental model to study oscillatory expression in a quiescent state.

**Figure 5:**
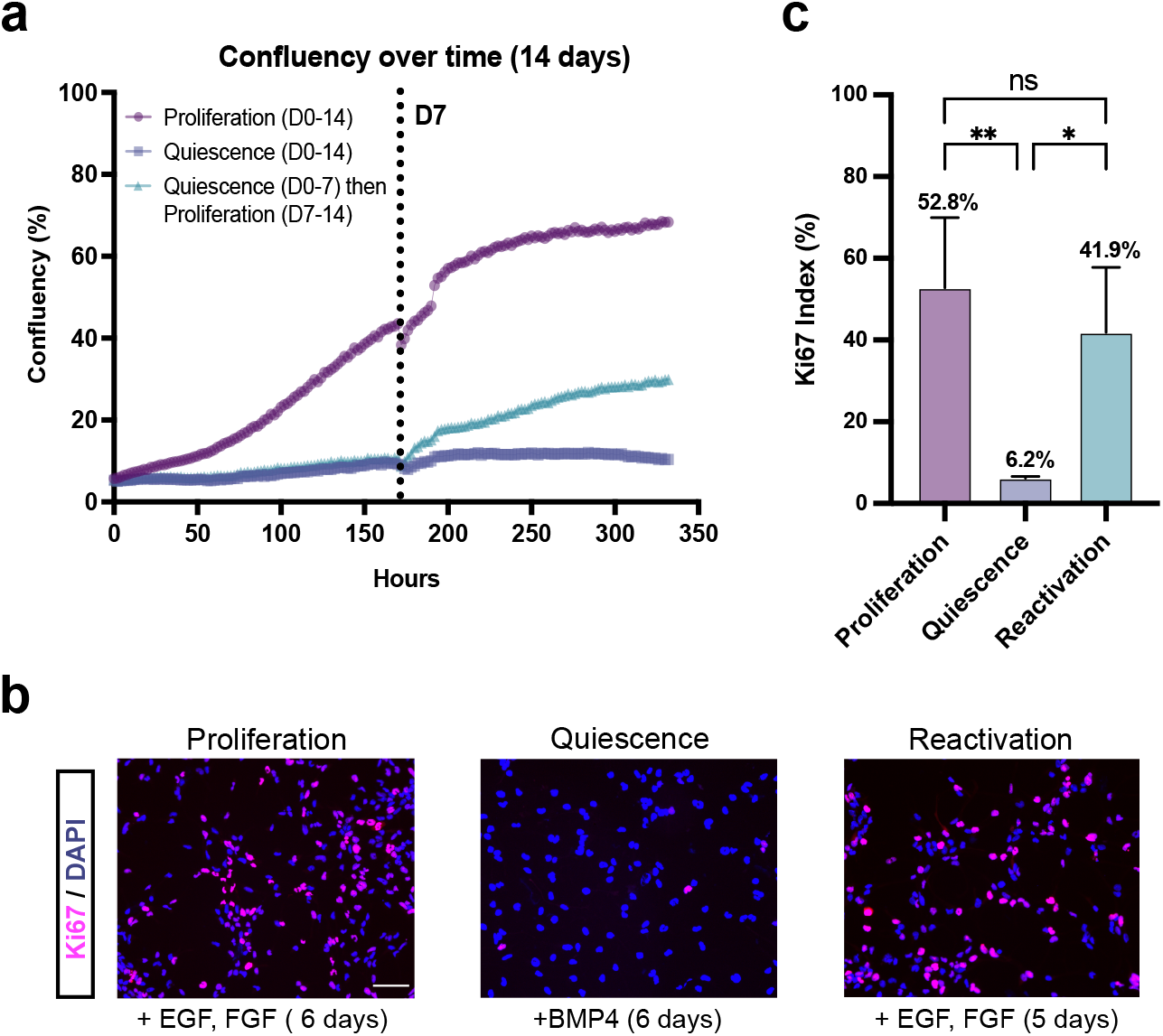
Inducing quiescence in GSCs. **(A)** Cell confluency analysis of GBM1 cells over a period of 14 days under 3 different conditions: Proliferation (cells cultured in proliferative media for 14 days), quiescence (cells cultured in quiescent media for 14 days) and reactivation (cells cultured in quiescent media for 7 days and then replaced by proliferative media for another 7 days). Vertical dotted line shows the time of media change for each condition with their respective media. **(B)** Immunofluorescence staining images of GBM1 cells for the proliferation marker Ki67 under 3 different conditions: proliferation (cells cultured in proliferative media for 6 days), quiescence (cells cultured in quiescent media for 6 days) and reactivation (cells were cultured in quiescent media for 6 days replaced by proliferative media for 5 days). Cells were also counterstained for DAPI (scale bar = 50 μm). **(C)** Bar graph showing the mean percentage of Ki67 positive cells under proliferative, quiescent and reactivated conditions (error bars represent standard deviation, n=3 biological experiments, total number of cells counted per condition Proliferation= 5448 cells, Quiescence= 3038 cells, Reactivation= 5828 cells, One-way ANOVA with Tukey’s multiple comparison test, proliferation vs quiescence **p=0.008, quiescence vs reactivation *p=0.021, proliferation vs reactivation ns= not significant).

### The stemness gene *SOX2* oscillates in proliferative and quiescent GSCs

From the core list of 4 candidate genes (*SOX2, JUNB, FOS, TSC22D1*) that have been predicted to oscillate and regulate the quiescent state (Fig. 4C), the stemness gene *SOX2* is of particular interest. A large number of studies in the literature have highlighted the important role of *SOX2* in promoting GBM malignancy ^39,40^ and the association of high SOX2 levels with tumour aggressiveness and poor clinical outcome ^41–43^. We therefore chose *SOX2* as our prime candidate to verify whether its protein expression oscillates in GSCs. To assess this we generated a *SOX2-mKate2* fusion knock-in line, using CRISPR/Cas9, to mark the endogenous SOX2 expression in GBM1 cells. In particular, we inserted a linker, followed by the mKate2 fluorophore and a P2A-Neomycin cassette, at the C-terminus of the SOX2 protein (Fig. 6A). This direct fusion enabled us to follow the kinetics of the endogenous SOX2 protein while maintaining the SOX2 protein intact. We generated 3 clonal *SOX2-mKate2* GBM1 lines which were all heterozygotes for *SOX2-mKate2*.

**Figure 6:**
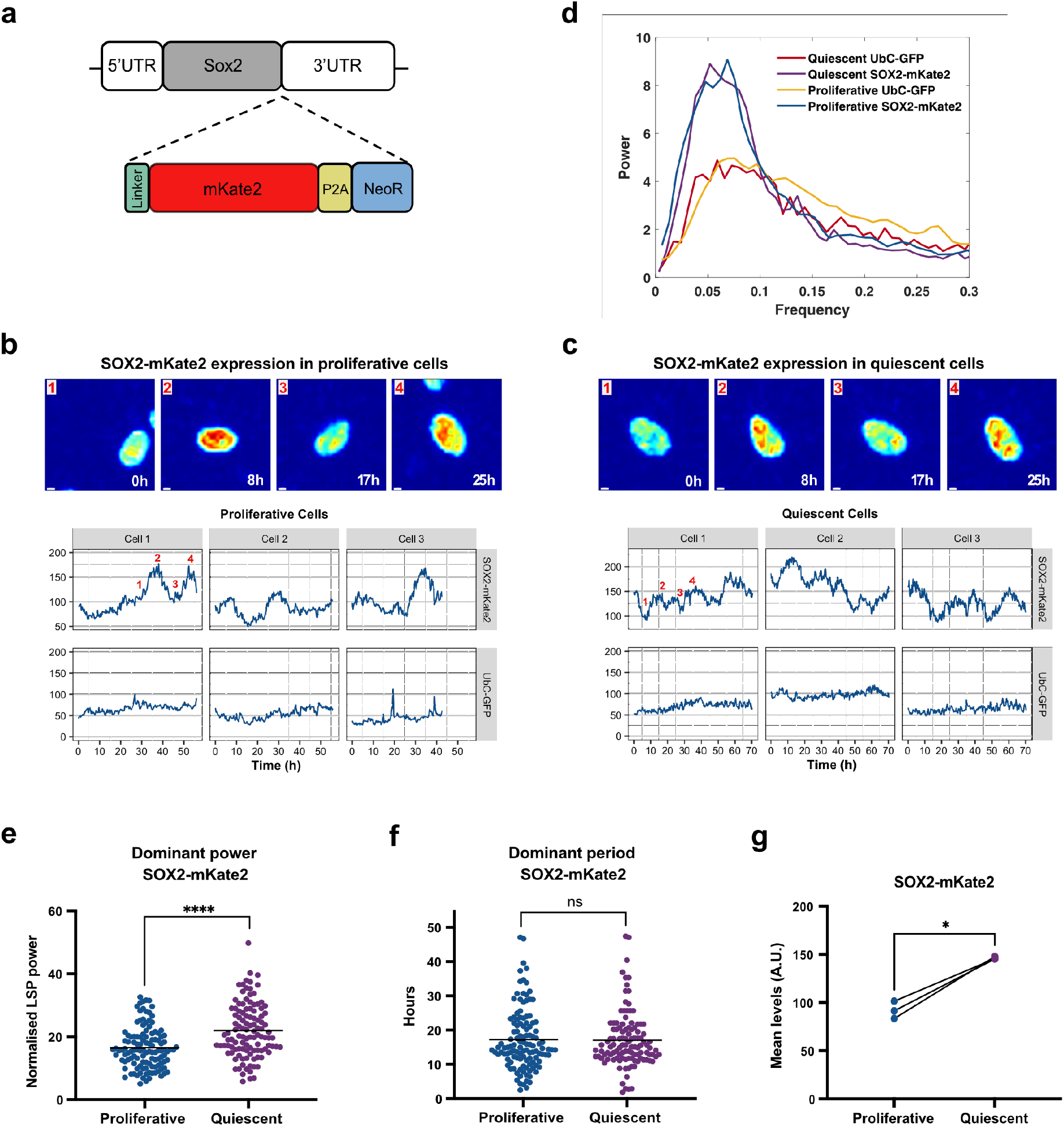
*SOX2* oscillates in proliferative and quiescent GSCs. **(A)** Schematic representation of the *SOX2* locus following genome editing with CRISPR/Cas9. A DNA sequence encoding for a linker protein, the mKate2 fluorescence protein followed by a P2A sequence and the neomycin resistance gene, has been inserted downstream of the *SOX2* exon and upstream of the 3’UTR. **(B-C)** Example snapshot images of SOX2-mKate2 expression in GBM1 cells in proliferative **(B)** and quiescent **(C)** conditions (top panels). Warm colours represent high expression and cold colours represent low expression (scale bar = 3 μm). Example cell traces showing SOX2-mKate2 and UbC-GFP expression over time in the same cell in GBM1 SOX2-mKate2 cells carrying a UbC-GFP reporter (A.U. = arbitrary units) (bottom panels). Numbers in Cell 1 in each condition indicate the timepoints that correspond to the images in the top panels. **(D)** Lomb-scargle periodogram (LSP) showing peaks in the power spectrum from SOX2-mKate2 and UbC-GFP expression in proliferative and quiescent conditions in GBM1 cells (averaged across all cell traces per protein expression per condition). SOX2-mKate2 expression shows a stronger peak in both proliferative and quiescent conditions compared to UbC-GFP **(E-G)** Graphs showing the dominant power **(E)** and dominant period **(F)** (as determined by LSP), and the mean protein expression levels for SOX2-mKate2 in proliferative and quiescent conditions (in (E) and (F) black horizontal lines represent mean, Mann-Whitney test, two-tailed, p****<0.0001, ns=not significant, in (G) dots represent mean levels per experiment, paired t-test, two-tailed, p*=0.012, n= 3 biological experiments, total number of cells tracked: Proliferative = 119, Quiescent = 112)

Next, we performed single-cell live fluorescence imaging in proliferative (Fig. 6B) and quiescent conditions (Fig. 6C), in GBM1 *SOX2-mKate2* cells. Tracking of the SOX2-mKate2 protein expression in single cells for approximately 50-70 hrs, revealed that SOX2 is dynamically expressed, showing peaks and troughs in intensity over time in single nuclei, where SOX2 is expressed (Fig. 6B-C). As a negative control, we tracked expression of the GFP fluorophore driven by the UbC promoter in the same cells, which had been previously transduced in the GBM1 cell line. Conversely to SOX2-mKate2, GFP expression showed no apparent or regular dynamicity, with GFP intensity forming a flat line over time denoting steady expression (Fig. 6B-C). To determine whether the SOX2-mKate2 protein expression oscillates, we performed periodicity analysis on multiple single cell intensity tracks using the Lomb-Scargle periodogram (LSP) ^44^ (Fig. 6D). Our analysis showed that the SOX2-mKate2 cell traces, in both proliferative and quiescent conditions, had a higher dominant power in the power spectrum compared to GFP traces, suggesting a stronger periodic signal (Fig. 6D). Further comparison of the SOX2-mKate2 dominant power between the proliferative and quiescent conditions revealed a significant increase in quiescent condition indicating that SOX2 oscillations are of better quality in the quiescent state (Fig. 6E). The average period of SOX2-mKate2 expression was calculated to be ∼17h in both proliferative and quiescent conditions (Fig. 6F), while the overall SOX2 levels (but not GFP (Supplementary Fig. S1)) were significantly increased by 1.6x fold in quiescence (Fig. 6G).

Overall, our findings show that SOX2 oscillates with the same period in both proliferative and quiescent conditions, suggesting that although the cells have entered quiescence, gene expression dynamics are maintained.

## Discussion

Gene oscillations are emerging as a powerful, yet largely unexplored, way to regulate gene expression and dictate cell-state transitions ^10,12,13,45,46^. In this paper, we have used a tailored bioinformatic pipeline ^21^ to identify genes that are expressed in an oscillatory manner in human GBM tumour samples. Based on our previous work, we have reasoned that such dynamically expressed genes are likely to be important for the transition of cells between states, such as a proliferative and a quiescent state ^10^. Here, we have identified a list of oscillators that are likely to control cancer progression in GBM and we have experimentally verified the stemness gene *SOX2* to oscillate in both proliferative and quiescent GSCs *in vitro*.

Genes expressed in an oscillatory manner are difficult to identify in static scRNA-seq data with most commonly used current bioinformatic methods. Time-series scRNA-seq or pseudotime inference methods fail to detect ultradian oscillations, which unlike circadian oscillations, tend to be out of phase between cells and therefore are very difficult to distinguish from expression noise. To get around this limitation, the approach that we used is based on pairwise gene comparisons, which looks at the co-expression relationship within each cell and across all cells and how well that fits a sinusoidal process, irrespective of real time information and accounting for a phase shift ^20,21^. However, two caveats remain: First, the absence of real time information precludes any direct estimation of periodicity. Second, our method detects potential oscillations at the mRNA level, which would be translated to protein expression oscillations only in appropriate production/degradation parameters ^47^. Indeed, in some cases mRNA oscillations were translated in a stepwise accumulation of the protein ^48^. Therefore, further experimental validation of the potential oscillations is needed, ideally by tagging the endogenous protein, as we have done for the case of *SOX2* here.

In this work, several steps have been taken to increase confidence in detecting potential oscillations. First, we verified that the inference pipeline can detect known oscillators such as HES1, Olig2 and ASCL1 ^11^ and cell cycle genes ^27^. Second, we chose genes that were consistently predicted to oscillate in each of the 5 tumour samples analysed (“Shared oscillators”) thus, less likely to be false positives. Third, we found that all shared oscillatory TFs (100%) were involved in gene network motifs capable of driving dynamic expression ^19^. Collectively these steps suggested that genes that were predicted to oscillate in our pipeline were high confidence candidates.

Our analysis predicted that approximately ∼3K genes (excluding cell cycle regulators) are likely to oscillate at the transcriptional (mRNA) level across all the tumours. Given that the number of genes previously shown to oscillate at the protein level across biological contexts is in the tens for the ultradian scale ^45,49,50^ and in the hundreds for the circadian scale (GO_0007623), rather than the thousands, this is a significant increase, suggesting that oscillations are a more widespread form of gene regulation that one might anticipate.

Interestingly, the “Shared oscillators” were enriched for functions that contribute to cancer pathogenesis, as opposed to “Non-oscillators”, which associated primarily with cell maintenance processes. In that regard, we hypothesise that oscillations are likely to enable cancer cells to develop adaptative strategies to survive and grow. Such examples we find in bacteria where studies have shown that microbial cells are known to use pulsatile expression to create a range of phenotypes that help them adapt a bet-hedging strategy to respond to environmental changes ^51^. Thus, our inference analysis can potentially help us identify key regulators that control cell-state transitions.

In GBM, quiescent GSCs have been found to be resistant to chemotherapy and radiotherapy ^52^ and responsible for re-initiating tumour growth following treatment ^33^. We have therefore looked specifically for oscillators that may control the quiescent state and identified a core list of 4 genes. From those, we selected the *SOX2* gene for further experimental validation. *SOX2* is well known for its role in maintaining pluripotency ^53^. In the context of GBM, *SOX2* has been implicated in promoting GBM malignancy ^39,43,54,55^. In addition, it was found that chromatin regions enriched for SOX binding motifs fail to reconfigure in response to BMP4, suggesting that the levels and/or activity of SOX proteins may impede the exit from self-renewal ^9^.

By generating an endogenous knock-in reporter we have found that *SOX2* oscillations are detected at the protein level in both proliferative and quiescent GBMs with a periodicity of 17hrs. This is a novel aspect of SOX2 expression that provides new insights into the way it may regulate downstream targets, whereby not just the presence or absence of SOX2 protein, but also its pattern of expression may dictate diverse cell-fate decisions.

Our findings are significant because dynamic gene expression in general, and ultradian oscillations in particular, have been linked to the ability of the cell to make cell-state transitions ^10,11,13,14,45,46^. SOX2 protein oscillations in GBM quiescent cells are indeed reminiscent of HES1 oscillations in quiescent neural stem cells. In the case of HES1, we and others have shown, that oscillations enable the exit from quiescence ^10,12^. It remains to be seen whether inhibiting SOX2 oscillations, but not affecting its overall expression, will prevent exit from quiescence of GBM cells. Locking cells in quiescence is an important therapeutic avenue to explore in addition to tumour resection and chemotherapy/radiotherapy. Alternative therapy strategies, such as promoting BMP4-induced tumour differentiation have been proven challenging with several barriers to implementation ^56^, as tumour cells fail to commit to differentiation and are susceptible to de-differentiation ^9^. On the other hand, silencing of *SOX2* showed promising results on reducing tumorigenicity ^57,58^, however abolishing SOX2 expression may have potential consequences in the normal brain, which is not a desirable outcome. Thus, understanding how SOX2 oscillations regulate the GBM quiescent state would be of great putative translational potential.

In summary, we have shown that oscillatory gene expression is prevalent in GBM neoplastic cells and characterises genes that are likely to play key roles in cell-state transitions. Deciphering this type of gene regulation, could have important therapeutic implications.

## Methods

### Identification of the neoplastic population

Single-cell sequencing data from 5 tumours (MGH 124, MGH 125, MGH 143, MGH 102 and MGH 115) in GEO GSE131928 (sample GSM3828673) ^22^ were used. All selected tumours were IDH-wild-type. Quality control checks and TPM (transcripts per million) calculations had already been undertaken by the study’s authors. For the identification of the neoplastic population, the integration method “Label Transfer” was used ^23^. The neuronal, vascular, glial, immune (GSE67835 ^59^) and neoplastic (GSE84465 ^24^) labels were used as a reference dataset and the number of anchors was set at 100. This method relies on similarities in gene expression programs between a reference and query dataset. The algorithm uses canonical correlation analysis and projects the datasets into a shared subspace defined by their correlation structure. The algorithm then identifies pairs of mutual nearest neighbours within the query and reference cells and these cells are used as “anchors” between reference and query datasets. This approach allows the labelling of overlapping cell types between reference and query dataset as well as the identification of unique cell types. In order to attain more homogenous subclusters of neoplastic cells, clustering was repeated on only the neoplastic population of each tumour using the Seurat (4.0.3.) package ^60^ with default parameters. These neoplastic subclusters were used as input for the OscoNet pipeline.

### Identification of low-cycling neoplastic subclusters

The ‘FindAllMarkers’ Seurat function ^60^ was used to determine differentially expressed genes (DEG) for each neoplastic subcluster in every tumour, with parameters ‘min.pct’= 0.25 and ‘logfc.threshold’ = 0.25. Only DEGs with an adjusted p value < 0.05 were further selected for Gene Set Enrichment Analysis (GSEA). GSEA was performed using the clusterProfiler package in R ^61^. The ‘msigdbr’ function was used to extract the human Hallmark gene sets from the MSigDB ^31^, the ‘pvalueCutoff’ was set at 0.1 and the ‘set.seed’ function was set at an arbitrary value of 1234. The results of this analysis are summarised in Supplementary Table S1. Clusters where one or more of cell-cycle related Hallmark gene sets (E2F Targets, G2M Checkpoint, Mitotic Spindle) were within the top 3 significantly enriched and downregulated gene sets were classed as ‘low-cycling’.

### Inference of oscillatory expression using OscoNet

Identification of oscillatory gene expression was performed for each neoplastic subcluster separately, using a modified version of the computational algorithm OscoNet ^21^. Briefly, for each neoplastic subcluster per tumour, we selected only genes that were expressed in at least 80% of the cells in each subcluster, and were highly variable (i.e. gene variance higher than the gene mean variance of the population). The algorithm then detected all potential pairs of co-oscillating genes and clustered them in communities with similar frequencies. Communities with just a single gene were excluded whereas all genes in the rest of the communities (regardless of the community significance score and linearity prediction status) were considered to be the inferred oscillators.

### Enrichment analysis

Data were analysed using the Qiagen Ingenuity Pathway Analysis ^28^ (QIAGEN Inc., https://digitalinsights.qiagen.com/IPA) ^62^ to assess enrichment for biological and molecular functions. The ShinyGO 0.77 tool (http://bioinformatics.sdstate.edu/go/) ^63^ was used with default package parameters to estimate enrichment for MSigDB gene sets in Fig. 3. As background list we used the list of all expressed genes (i.e. all genes passed zero-filtering per neoplastic subcluster per tumour) (Supplementary Table S10).

### Dynamic gene network motif analysis

We employed a previously published bioinformatic screen to identify TFs involved in dynamic gene networks, which consist of an autoregulatory TF that co-regulates a miRNA ^19^. The screen was performed by integrating published TF binding site (TFBS) from the ReMap 2022 database ^64^ and miRNA target data from miRbase (Release 22.1) ^65^ and miRTarBase (release 9.0) ^66^ into a single network. This network was then analysed to reveal genes involved in dynamic gene network motifs. For this work, the original network was re-built and modified to include the latest available TFBS in ReMap and miRNA databases.

### Cell Culture

The GBM1 cell line was derived from IDH-wild-type primary GBM tumours as previously described ^36^ and was kindly donated by the Wurdak Lab (Stem Cell and Brain Tumour Group, University of Leeds). The line had been further modified to carry a UbC promoter driven GFP fluorophore reporter (GenTarget Inc, LVP1229), via viral transduction as previously described ^37^. Puromycin (2 μg/ml) was used to select for positive cells for UbC-GFP expression. Cells were cultured on L-Laminin and Poly-L ornithine pre-coated flasks/plates in the presence of Neurobasal Medium (Gibco, Cat# 21103-049) supplemented with 0.5% N2 (100x) Supplement (Life Tech, Cat# 17502048), 1% B27 (50x) Supplement (Life Tech, Cat# 17504044), FGF (40 ng/ml) (PeproTech, Cat# 100-18B) and EGF (40 ng/ml) (PeproTech, Cat# 315-09) (proliferating media). For coating, any flasks/plates were first coated with Poly-L ornithine (Gibco, Cat# P3655) at 5 μg/ml in water for 1 h at 37oC, followed by 1x wash with water and coated subsequently with L-Laminin (Sigma-Aldrich, Cat# L2020) at 2 μg/ml in PBS overnight at RT, followed by 1x wash with PBS before cell seeding. For induction of quiescence, the FGF and EGF mitogens in the media were replaced by 100 ng/ml of BMP4 (Recombinant human BMP-4, PeproTech, Cat# AF-120-05ET) (quiescence media). To reactivate cells from quiescence, quiescence media was replaced by proliferating media after a gentle 2x wash with PBS.

### Generation of SOX2-mkate2 fusion cell line

We used CRISPR/Cas9 to generate a C-terminally endogenously tagged SOX2-mKate2 protein fusion in the GBM1 cell line. The donor template for Homologous Directed Repair (HDR) containing the Glycine-Alanine linker-mKate2 fluorophore-P2A-neomycin resistance (NeoR) cassette was PCR-amplified from the eFlut Plasmid 1A collection for C-terminus fluorescence tagging, kindly donated by the Lahav Lab ^16^. The primers used to amplify the HDR template had 40bp homology to the *SOX2* gene and 20 bp homology to the template and are listed in Supplementary Table S11. Guide RNAs (gRNAs) targeting the human *SOX2* stop codon sequence were designed using the CRISPR Finder tool on the Wellcome Sanger Institute Genome Editing (WGE) website (https://wge.stemcell.sanger.ac.uk//, 2021) and are listed in Supplementary Table S11. gRNAs were obtained as crRNAs and were mixed with tracrRNA in equal ratios of 45 μM in IDT duplex buffer (IDT, Cat# 1072570), heated at 95°C for 5 minutes and then cooled slowly to RT. The Cas9 was delivered in the form of Ribonucleoprotein (RNP) complex, where the annealed gRNA-tracrRNA were complexed with the Cas9 protein (Alt-R^TM^ S.p. HiFi Cas9 Nuclease V3, IDT, Cat# 1081061) in IDT duplex buffer at RT for 10 minutes before adding 1ug of the donor template (final concentrations: each gRNA 20.4 μM, Cas9 protein 11.09 μM, donor 9 ng/μl). The RNP complex was mixed with 8 x 10e5 GBM1 cells resuspended in 100 μl of SG solution from the SG Cell Line 4D-Nucleofector^TM^ X Kit (Lonza, Cat# V4XC-3024). This entire mix was then placed in a Nucleocuvette^TM^ and was nucleofected using a Lonza Amaxa 4D-Nucleofector ^TM^ and the EN138 nucleofection programme. Following nucleofection, cells were further cultured in the presence of 200-400 μg/ml Neomycin (G418 Sulfate, ThermoFisher Scientific, Cat# 11811023) to select for genetically modified cells. Antibiotic resistant cells were then subjected to single-cell FACS for mKate2 and GFP expression (double positive) using a BD Influx sorter and plated onto pre-coated glass (Greiner BIO-ONE Sensoplate^TM^ glass bottom, Cat# 655892) or plastic (Merk Corning^©^ TC-Treated Microplates, Cat# CLS3997) bottom 96 well plates. An average number of 7 clones per 96-well plate grew into clonal lines and where further expanded. 10 clonal lines in total were screened by PCR for incorporation of the donor sequence and 9 were found to be positive. The genotyping primers used are listed in Supplementary Table S11. All positive clones were HET for the knock-in fusion. 6 clonal lines were further submitted for Sanger sequencing which verified that 5 out of 6 clonal lines had an intact wild type allele and they had correctly integrated the donor sequence. 3 out of those 5 clonal lines were further selected for experimentation.

### Immunostaining

GBM1 cells were cultured in Poly-L ornithine and L-Laminin pre-coated coverslips in either proliferating media, quiescent media (for 6 days) or in proliferating media after reactivation from quiescence (that is 6 days in quiescent media followed by 5 days in proliferation media). Immunostaining was performed as previously described ^10^. Primary antibody was mouse anti-Ki67 (1:200, BD Biosciences, Cat #550609) and secondary antibody was anti-mouse 568 (1:500, Thermo Fisher Scientific, Cat#A-11004). Coverslips were mounted with ProLong Gold Antifade mountant with DAPI (Invitrogen, Cat#P36935).

### Image and time-lapse movie analysis

Confluency analysis of GBM1 UbC-GFP cells was performed using the ZOOM Incucyte Live-Cell imaging platform (Sartorius). 8K cells per square centimetre per well were plated in 6-well plates and whole-well imaging was performed in 120-minute intervals using a Nikon 4x objective. Cells were cultured continuously either in proliferative or quiescence media. For reactivation, quiescence media was replaced by proliferative media after 7 days in quiescence. Images were processed using the IncuCyte® ZOOM 2018A software to determine cell confluency over time.

Immunofluorescence images were captured using a Nikon Eclipse 80i with a Plan Fluor 20x / 0.5 DIC N2 objective and a 1280 x 1024 format. Image processing of Ki67/DAPI staining, to quantify number of Ki67 positive nuclei, was performed on ImageJ 2.1.0/1.53c using automated tools for detection of fluorescence expression. Immunostaining images with just the secondary antibody were used as controls to determine the expression detection threshold.

For dual fluorescence imaging of GFP and mKate2, GBM1 SOX2-mKate2 clonal cells carrying the UbC-GFP reporter, were plated in a 4-compartment pre-coated glass bottom 35 mm dish (Greiner BIO-ONE, Cat# 627870). 22K cells per square centimetre were plated in one compartment and imaged after being kept in proliferating media for 7 days and 44K cells per square centimetre were plated in another compartment and imaged at the same time after being kept in quiescence media for 6 days (proliferation media was replaced by quiescence media 1 day post cell seeding). Time-lapse imaging was performed for a total of 70-90 h using a Zeiss LSM 880 Inverted Airyscan with a Plan-Apochromat 20x / 0.8 M27 and a 1536 x 1536 or 1536 x 1024 format. Images were acquired every 20 min with z-stacking covering a 35.2 um depth. The fluorescent signals were collected on different tracks. For detection of GFP and mKate2 a 488 nm and a 594 nm laser were used for excitation and the emitted fluorescence was collected between 490-552 nm and 597-695 nm respectively. The fluorescent images were analysed using the Imaris imaging software (Bitplane) and individual cells were tracked using the Imaris “Spot” function to quantify the mKate2 and GFP fluorescence intensity signal over time.

### Periodicity analysis

To estimate periodicity of the traces of SOX2-mKate2 expression we developed a MATLAB pipeline which made use of the Lomb-Scargle periodogram. This is an algorithm for detecting and characterizing periodic signals in unevenly sampled data^44^.

The steps used to develop our estimates were as follows: firstly, we detected a trend in each trace via a Gaussian-weighted average filter with a 24 hr moving window, as we were focusing on fluctuations that have less than 24 hr periodicity. We then subtracted this trend line from the raw data to obtain a new detrended trace, which was then fed into the “plomb” function that outputs the Lomb-Scargle power spectral density (PSD) estimate, sampled at the instants specified in an associated frequency vector.

The position of a spike in the X direction of this PSD estimate corresponds to the dominant frequency (1/period) at which the detrended data trace is oscillating. Conversely, more low and shallow outputs in the PSD suggests a lack of oscillatory behaviour in the trace. To provide an average periodicity estimate we used the “interp1” function to interpolate the different traces, such that time points within each trace were consistent and therefore comparable. This allowed for a mean PSD/frequency plot to be developed for both the proliferative and quiescent conditions. This was also done on GFP traces as a control. Cell traces with time duration shorter than 25 h and expression variance higher than 5 x standard deviations from the mean were excluded. The excluded data accounted for 4% of overall data acquired in proliferation and less than 2% in quiescence conditions.

### Statistical analysis

Statistical analysis was performed using GraphPad Prism 10.0.3. A Shapiro-Wilk normality test was undertaken to determine whether a parametric or non-parametric statistical test should be used. When sample size was too small (i.e. n=3), the distribution was judged based on the Normal QQ plots. The alpha level was set to .05.

The statistical tests undertaken, the number of biological replicates, the sample size and p-values are reported in the relevant figure legends.

### Data availability

Raw imaging data and processed datasets have been deposited at Figshare and will be made publicly available upon publication. Any data reported in the current study that are not present in the public repository, are available from the corresponding author on reasonable request.

## Additional Information

### Acknowledgements

The authors wish to thank Prof. Andrew King and Dr David Coope for their mentorship, Prof. Steve Pollard, Prof. Stuart Allan and Prof. Omar Pathmanaban and Assoc. Prof. Ryan Mathew for advice and discussions, Dr Bronwyn Irving, Christopher Akhunbay-Fudge and Holly Briggs for assisting with cell culture, Alexander Landry for assisting with bioinformatic analysis and Alessandro Ricci for his help with optimizing and scaling the label transfer technique, Dr Mike Croucher for assisting with implementation of OscoNet and all members of the Papalopulu lab and the Surgical Neuro-Oncology Manchester lab for comments on the work. The authors would also like to thank the Bioimaging Facility (especially Dr Peter March and Dr David Spiller for their help with imaging) and the Flow Cytometry facilities (especially Dr Gareth Howell) of the University of Manchester for technical support. The Bioimaging Facility microscopes used in this study were purchased with grants from BBSRC, Wellcome Trust and the University of Manchester Strategic Fund. This work was supported by a Wellcome 4Ward North Clinical PhD Fellowship to R.Z.F. (203914/Z/16/Z), a Sir Henry Wellcome Trust Postdoctoral Fellowship to E.M (201380/Z/16/Z) and a Wellcome Trust Senior Research Fellowship to NP (224394/Z/21/Z).

### Author contributions

R.Z.F. and E.M. performed experiments and data analysis. O.C., L.C., A.R. and Z.Z. performed data analysis and contributed to writing the relevant sections in the manuscript. E.M. and N.P. supervised the study. H.W. co-supervised the study and provided experimental material. R.Z.F., E.M. and N.P. conceived the study. E.M. and N.P. co-wrote the manuscript with input from R.Z.F. All authors gave their approval for publication.

