## Supplementary Information for "Identification of genes with oscillatory expression in glioblastoma – The paradigm of *SOX2*"

#### Supplementary Fig. S1

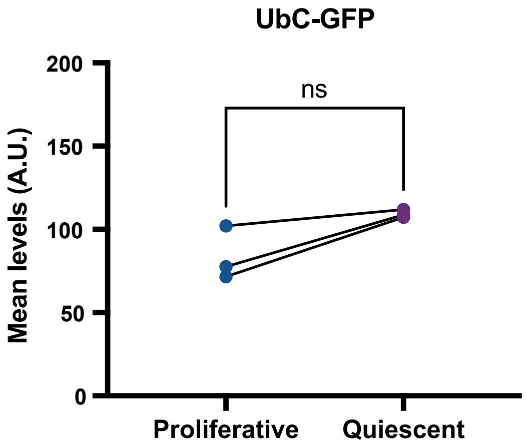


**Supplementary Fig. S1: Comparison of UbC-GFP expression levels in proliferative vs quiescent conditions**
Graph showing the mean protein expression levels for GFP in proliferative and quiescent conditions, dots represent mean levels per experiment, paired t-test, two-tailed, ns = not significant, n=3 biological replicates, total number of cells tracked: Proliferative = 119, Quiescent = 112

### Supplementary Methods

#### Assessing the effect of zero gene transcription expression on the OscoNet performance

We simulated a synthetic experiment with 100 single cells and 1000 genes, of which 400 were samples from a sinusoidal function with additive Gaussian noise. The 400 oscillators were simulated in 2 frequency groups, each group containing 200 genes. The relative frequencies of the two groups were proportional to 2:3.

In each group, half of the oscillatory genes were simulated as strong oscillators with noise variance $\sigma^2$. The other half were simulated as weak oscillators with noise variance (2σ)^2. The starting phase $\Phi$ varies in different genes within a frequency group. The remaining genes were simulated as independent Gaussian noise.

To explore the effect of zero gene transcription expression in the overall downstream analysis we varied the percentage of zeros (sparsity) from 0 to 1 with step 0.05, in the expression level of half of the genes in the 100 cells for each of the frequency groups.

For each level of sparsity, we run the OscoNet pipeline and then computed the True Positive Rate (TPR), False Positive Rate (FPR) and False Discovery Rate (FDR) defined as follows and depicted in Supplementary Table S12

TP= number of true co-oscillating pairs
TN= number of true non co-oscillating pairs
FP= number of false co-oscillating pairs
FN= number of false non co-oscillating pairs
FPR= FP/Actual Negative= FP/(FP+TN)
TPR=TP/Actual Positive = TP/(TP+FN)
FDR =FP/ Predicted Positive = FP / (FP + TP)

Increasing the percentage of zeros in the true co-oscillators does not affect the FPR or the FDR that stays close to the suggested threshold 0.05 (Supplementary Fig. S2) Hence, the presence of zeros in our simulated examples does not introduce bias towards the positive discoveries (i.e. number of genes incorrectly identified as co-oscillators). At the same time, we note that an increase of sparsity affects the capability of detecting co-oscillating genes. Indeed, when the sparsity increases from 0 to 0.2, the FPR suffers a fast decrease from 1 to 0.54 while for sparsity greater than 0.2 the FPR falls below 50% (Supplementary Fig. S2). This means that in the pessimistic scenario for which half of the co-oscillating genes in our experiment were affected by at most 20% sparsity, we expect to identify at least 50% of the co-oscillating genes, with an FDR smaller than 0.05. Hence, we suggest 0.2 as sparsity threshold.

| | Actual Positive | Actual Negative |
|---|---|---|
| Predicted Positive | TP | FP |
| Predicted Negative | FN | TN |

**Supplementary Table S12: Confusion matrix representing the intersections between predicted and actual values in terms of TP, TN, FP and FN.**

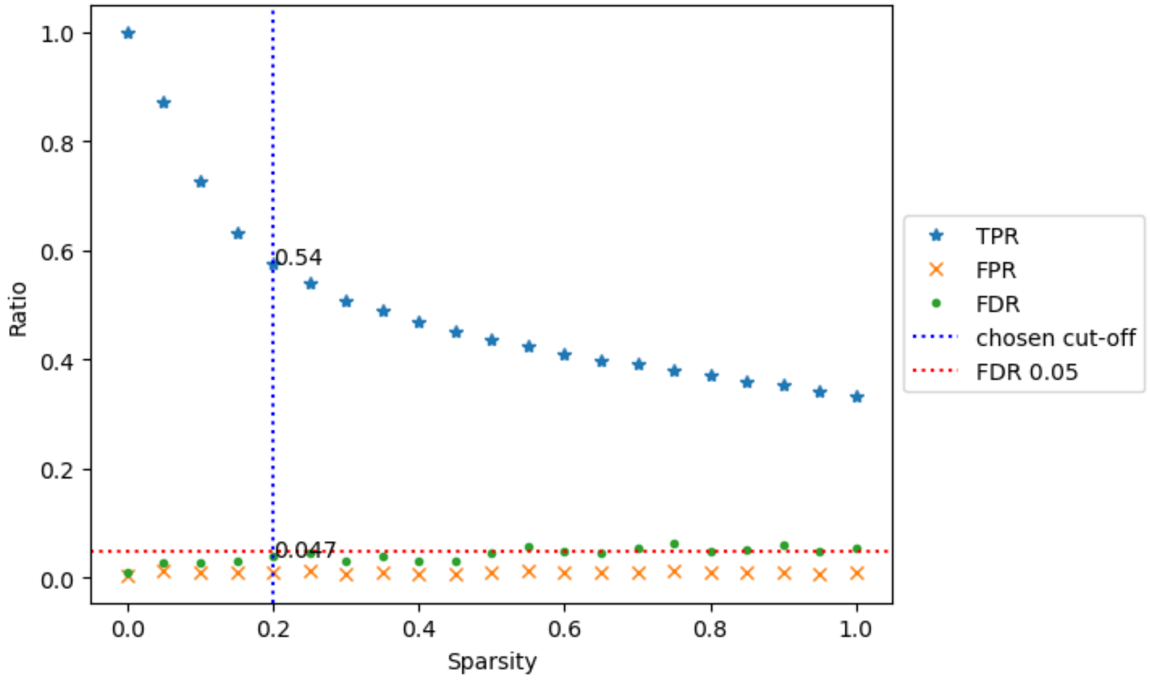


**Supplementary Fig. S2: Estimating True Positive Rate, False Positive Rate and False Discovery Rate**

Graph showing the TPR, FPR and FDR for every simulated set of data at different level of sparsity [0,…,1]. The vertical dashed blue line represents the suggested cut-off threshold for the maximum number of zeros for a gene to be retained in our analysis. The horizontal red dashed line represents the ideal FRD threshold of 0.05.
